# Identifying Core Operons in Metagenomic Data

**DOI:** 10.1101/2019.12.20.885269

**Authors:** Xiao Hu, Iddo Friedberg

## Abstract

An operon is a functional unit of DNA whose genes are co-transcribed on polycistronic mRNA, in a co-regulated fashion. Operons are a powerful mechanism of introducing functional complexity in bacteria, and are therefore of interest in microbial genetics, physiology, biochemistry, and evolution. Here we present a Pipeline for Operon Exploration in Metagenomes or POEM. At the heart of POEM lies the concept of a core operon, a functional unit enabled by a predicted operon in a metagenome. Using a series of benchmarks, we show the high accuracy of POEM, and demonstrate its use on a human gut metagenome sample. We conclude that POEM is a useful tool for analyzing metagenomes beyond the genomic level, and for identifying multi-gene functionalities and possible neofunctionalization in metagenomes. Availability: https://github.com/Rinoahu/POEM_py3k

## Background

It is estimated that 5-50% of bacterial genes reside in operons [6, 44], and the characterization and understanding of operons is central to bacterial genomic studies. Experimental approaches, chiefly RNA-Seq, are the most reliable way to identify operons; however, it is not feasible to perform experiments to characterize all operons. Over the years, several computational operon-prediction techniques have been developed. Generally, computational operon identification methods include three steps: 1. identify genes that are in an operon and, conversely, genes that do not participate in an operon; 2. identify features typical of each group; 3. train a classifier with these features and build a discriminating model.

Computational operon prediction methods have been developed since the late 1990’s (For a comprehensive review see: [46]). Naïve Bayes models have been used since early 2000’s for predicting operons [3, 10, 18]. Another method used microarray data to identify the different expression profiles of adjacent gene pairs in operons and outside of operons. The differential expression profiles and intergenic distances were used as as features to train a Bayesian classifier [35]. Comparative genomic methods were also used to identify operons by detecting conserved gene clusters across several species [5, 26, 31]. Other methods include particle swarm optimization [8, 9], and neural networks [39].

There are several operon databases that include automated and experimental-based operon annotation [13,25,29,33,38]. However, a manual curation method is not suitable for the rapid growing number of bacterial genomes, few of which are experimentally assayed for operons. Furthermore, experimental studies tend to use data from model species, and cross-species prediction may not work well [11].

The challenge of discovering operons is compounded when trying to discover operons in metagenomic data. Major additional confounders include the large loss of genomic information, short contigs that rarely assemble into a full genome, and misassembly that might produce chimeric contigs [45]. At the same time, metagenomic data contain rich information that cannot be gleaned from clonal cultures; it is therefore necessary to investigate how well we can predict operons in metagenomic data. Some work has been done including use of proximity and guilt-by-association [41, 42].

While a genome contains the total genetic information of an organism, a metagenome is a partial snapshot of a population of genomes. We therefore can rarely expect an operon discovery method to provide the entire content of operons from metagenomic data. However, predicting whether genes participate in an operon, and which functions are carried out by operons, provide valuable additional information to the functional annotation of a metagenome. In this study we present a method that (1) classifies gene pairs in metagenomes into “operonic” and “non-operonic” classes, and (2) provides functional annotations for the operons it reconstructs from metagenomic data. We introduce the concept of metagenomic *core operons*. A core operon comprises a set of intra-operonic gene pairs that have orthologs in several species in the metagenome, and are concatenated using guilt-by-association. Additionally, we introduce the *core functions* of operons, which identifies which functions in the metagenome are executed by operons. Commonly, metagenomic analysis pipelines provide the distribution of biological function the metagenome has based on a normalized count of functionally-annotated ORFs. Our method, a <u>P</u>ipeline for <u>O</u>peron <u>E</u>xploration in <u>M</u>etagenomes or POEM, adds more information as it considers the evolutionary conservation of co-transcribed genes in the species constituting the microbial community. This additional information is valuable for understanding the genetic potential of a microbial community introducing structural information in the form of predicted operons.

## Results

We ran POEM on two different data sets. One includes simulated reads generated by ART [15] from 48 genomes of Operon DataBase v2 [29]. The genome species used and parameters of ART are shown in Supplementary Table S1. The second set is the human microbiome set SRR2155174 downloaded from ENA [21]. As a standard of truth for the operons, we used operons from Operon DataBase v2 that are supported by literature (henceforth: “true operons”). This dataset contains 8,194 genes and 5,621 adjacent genes in 2,589 operons.

### Metagenome Assembly and Gene Prediction

We used IDBA-UD, MegaHIT, and Velvet [47] to assemble the simulated and experimental reads; the results are shown in Table 1. IDBA-UD provided the maximal N50 and minimal number of contigs in both datasets. MegaHIT provided the largest genome size and the most protein-coding genes.

**Table 1.** Main features of simulated and real metagenome assembly. **Size (bp):** size of assemblies without singleton reads; **Genes from the True Operon Set**: genes discovered by the gene calling software, that are found in the True Operon Set (fraction of 8,194 found).

| Feature\Assembler | Simulated Reads |  |  | SRR2155174 |  |  |  |
| --- | --- | --- | --- | --- | --- | --- | --- |
|  | IDBA-UD | Megahit | Velvet.51 | IDBA-UD | Megahit | Velvet.51 | Assembly |
| Size (bp) | 132,218,137 | 134,341,573 | 100,889,182 | 131,424,989 | 135,150,882 | 85,261,552 | 122,371,235 |
| GC content | 49.51% | 49.55% | 50.73% | 48.11% | 48.19% | 46.96% | 48.04% |
| Number of contigs | 48,508 | 54,274 | 61,093 | 87,992 | 107,718 | 146,313 | 55,925 |
| Max contig length | 947,260 | 549,191 | 569,707 | 484,034 | 249,170 | 106,439 | 327,893 |
| Min contig length | 100 | 200 | 101 | 100 | 200 | 101 | 500 |
| Mean contig length | 2,725 | 2,475 | 1,651 | 1,493 | 1,254 | 582 | 2,188 |
| N50 | 12,732 | 7,681 | 9,312 | 4,306 | 2,593 | 906 | 4,331 |
| N90 | 984 | 888 | 569 | 478 | 426 | 227 | 754 |
| Protein-coding genes | 154,908 | 162,282 | 133,298 | 175,983 | 190,946 | 170,100 | 147,873 |
| Genes from the True Operon Set | 5,116 (0.62) | 5,078 (0.62) | 3,530 (0.43) | NA | NA | NA | NA |

Metagenemark found 7,855 genes of the 8,194 true operon genes in the whole genomes. In the simulated reads assembly, the number of genes numbers are 5,116, 5,078, and 3,530 (out of 8,194) using IDBA, Megahit, and Velvet respectively.

### Operon Prediction and Adjacent Genes Within the Operon

We tested the operon prediction module’s performance on whole genomes and simulated metagenome assembly. The 4,425 operonic and 2,097 non-operonic adjacent genes mentioned above were used as a True Positive (TP) set and True Negative (TN) set, respectively. The precision, recall, and F_1_ for predicted operonic adjacency are defined in the following equations and the statistical results are shown in Table 2.

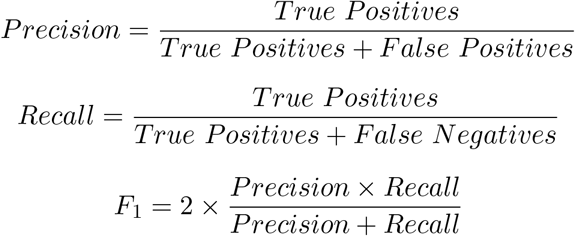

**Table 2.**
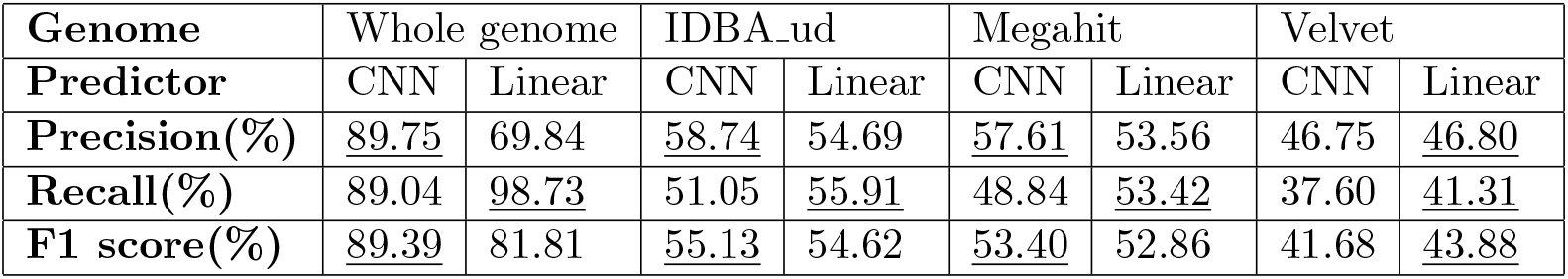
Evaluation of operonic adjacency prediction on simulated metagenomes. **CNN**: a convolutional neural network based classifier; **Linear**: a linear classifier that is based on the intergenic distance and strand co-location.

However, these results only reflect POEM’s performance on classification of operonic and non-operonic adjacency. To further evaluate POEM’s performance on full operon prediction, we report on the precision / recall analysis as illustrated in Figure 1. The total number of true operons in the simulated metagenome was determined to be 2,589. The results are shown in Table 3. POEM’s CNN performs much better than the linear baseline method when tasked with a perfect recovery of operons. For a 0.6 or better recovery, the CNN and the baseline perform similarly. This suggests that high quality longer assemblies, perhaps from longer reads, may perform better.

**Table 3.** Predicting operons in whole genome and assemblies of simulated metagenomes. ≥ **recovery**: ≥ 60% genes in a predicted operon belong to a known operon; **Perfect recovery**: both precision and recall equal one. Table shows number of operons recovered, and (fraction of 2,589)

| Real Operon | Whole genome |  | IDBA |  | Megahit |  | Velvet |  |
| --- | --- | --- | --- | --- | --- | --- | --- | --- |
| Classifier | CNN | Linear | CNN | Linear | CNN | Linear | CNN | Linear |
| **≥0.6 recovery** | 1,997 (0.77) | 1,816 (0.70) | 1,232 (0.48) | 1,322 (0.51) | 1,332 (0.51) | 1,377 (0.53) | 914 (0.35) | 881 (0.34) |
| **Perfect recovery** | 1,025 (0.40) | 469 (0.22) | 526 (0.20) | 253 (0.10) | 470 (0.18) | 233 (0.09) | 360 (0.14) | 160 (0.06) |

**Figure 1.**
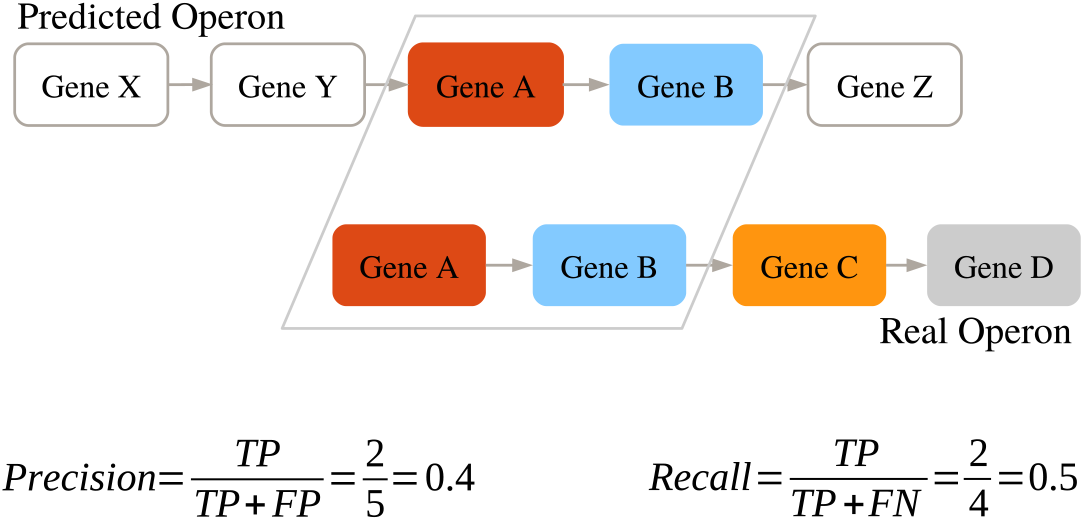
Determining precision and recall for a predicted operon.

### Core Functions Facilitated by Predicted Operons in Metagenomic Data

To functionally analyse operons in metagenomes we use core operons, which are described in the Background section. Briefly, core operons are weighted-edge undirected graphs that capture information about predicted orthologous operons or subsets of operons in the metagenome. The nature of the fragmented and partial nature of metagenomic data prohibits a clear binning of reads and a full assembly into component genomes. Therefore, we may not be able to provide an accurate prediction of all genes in the operons or their precise taxonomic affiliation. See Methods / Constructing Core Operons and Figure 5 for an explanation of how core operons are constructed. To see how well core operons capture the function of true operons on our different data sets, we examined the overlap of operonic genes with identical functions as shown in Figure 5. The results of this analysis is shown in Table 4.

**Table 4.** Comparing core operons discovered by POEM in the simulated metagenome, and in SRR2155174. See Methods and Figure 5 for details. **Intersection with True Operons**: The number of shared core functions between true operons and predicted operons. **SE**: standard error.

| Assembler | Number of core operons | Intersection with True Operons | Mean Precision ± SE | Mean Recall ± SE | Mean $F_1 \pm$ SE |
| --- | --- | --- | --- | --- | --- |
| True Operon Set |  |  |  |  |  |
| NA | 110 | NA | $0.97 \pm 0.12$ | $0.66 \pm 0.28$ | $0.75 \pm 0.22$ |
| Genome |  |  |  |  |  |
| NA | 310 | 36 | $0.77 \pm 0.34$ | $0.60 \pm 0.34$ | $0.67 \pm 0.28$ |
| Simulated Reads |  |  |  |  |  |
| IDBA_ud | 260 | 48 | $0.83 \pm 0.31$ | $0.61 \pm 0.33$ | $0.71 \pm 0.26$ |
| Megahit | 256 | 46 | $0.84 \pm 0.31$ | $0.61 \pm 0.32$ | $0.71 \pm 0.26$ |
| Velvet | 202 | 56 | $0.87 \pm 0.30$ | $0.59 \pm 0.32$ | $0.71 \pm 0.25$ |
| SRR2155174 |  |  |  |  |  |
| IDBA_ud | 141 | 25 | $0.71 \pm 0.36$ | $0.55 \pm 0.33$ | $0.65 \pm 0.24$ |
| Megahit | 138 | 26 | $0.71 \pm 0.37$ | $0.54 \pm 0.33$ | $0.66 \pm 0.24$ |
| Velvet | 94 | 11 | $0.72 \pm 0.39$ | $0.48 \pm 0.33$ | $0.65 \pm 0.25$ |

To show the utility of our method in discovering core functions facilitated by predicted operons, we ran POEM on the metagenome sample SRR2155174, containing the human gut microbiome data. Figure 2A shows a core function predicted from the SRR2155174 data set. The annotations of the core functions indicates that it is related to lipid transport and metabolism. We found several predicted operons (Figure 2B-E) from the SRR2155174 data set that match the core function. Of the loci in the core operon, only lp 1674 and lp 1675 loci in *Lactobacillus plantarum* WCFS1 (Figure 2E) can be found in the predicted operons of Operon DataBase [29]. To find the functions of these predicted operonic genes, we examined the functional annotations for these operonic genes from GenBank [2]. The functional annotations (Supplementary Table S2) show that these operonic genes are likely to be involved in fatty acid biosynthesis. We mapped the predicted operonic genes of *Lactobacillus plantarum* WCFS1 (Figure 2E) to KEGG database [19] and found most of the genes involved in fatty acid biosynthesis (Supplementary Figure S2). These results show these predicted operons are likely involved in fatty acid biosynthesis and have a high probability of being true operons. Although core operons are involved in the same biological pathway, the genes outside the core function (Figure 2B-E) are diverse. The core function reflects the conservation of operons across species and is more robust and error-tolerant than operons. Core functions may reconstruct the metabolism pathways from the incomplete genome assembly data leveraging the conservation of genes across species. The ability to use core functions as familiar ground from which to explore new conserved proximal genes makes core functions a new and powerful tool for discovering novel operon-encoded pathways in metagenomic data.

**Figure 2.**
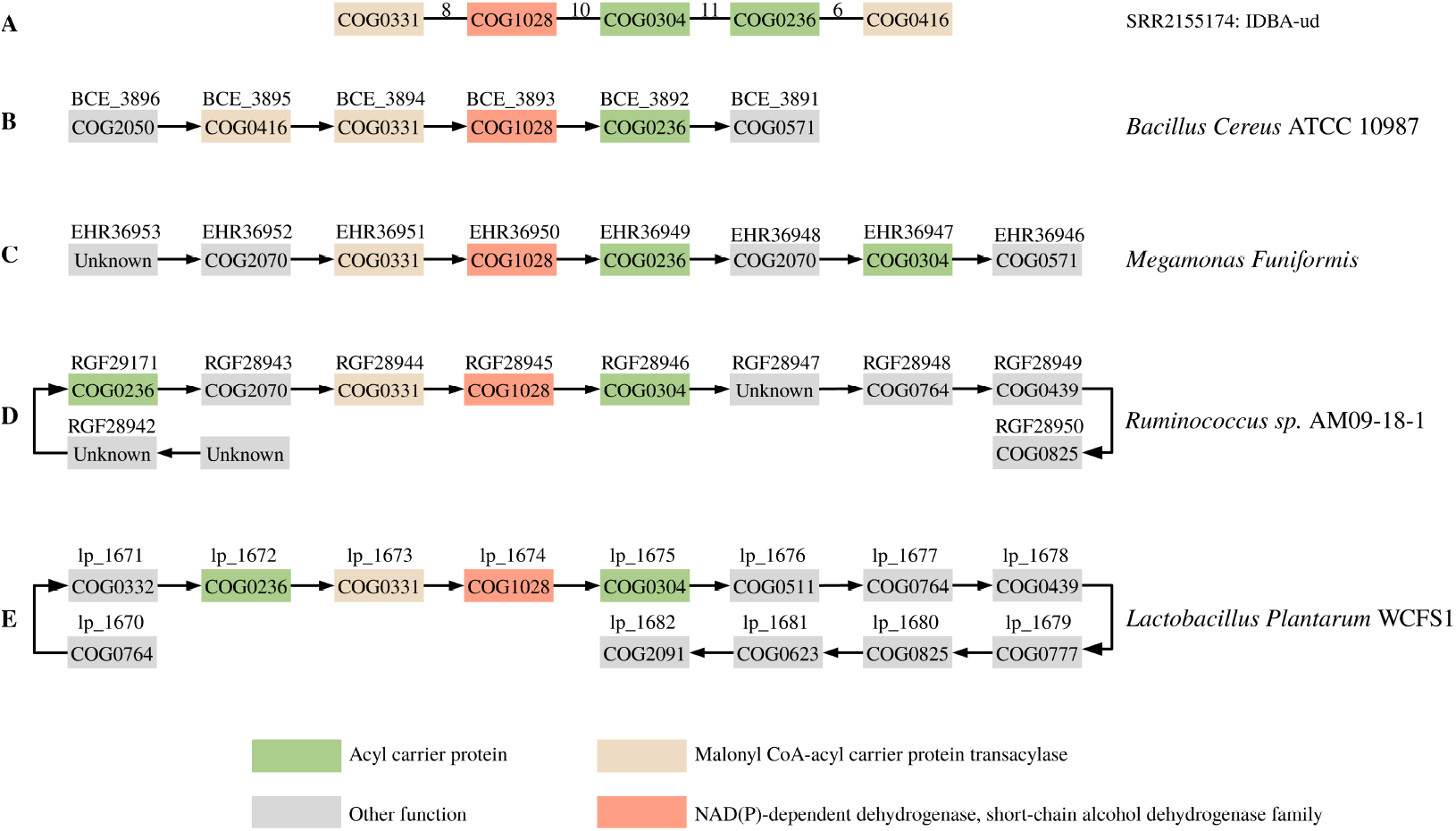
Mapping core functions to predicted operons. **A**: predicted core function from SRR2155174 data set; **B-E**: predicted operons in different species. the arrows stand for the strands of genes, box color is the COG functional classification; gray boxes are functions outside the core operon. Gene names are above the boxes.

## Discussion

In this study we introduce POEM, a complete pipeline for predicting operons in genomic and metagenomic data. We also introduce the concept of a core operon, a functional unit of proximal genes in a metagenome, which is composed of the common functions of orthologous operons. POEM’s CNN predicts intra-operonic genes with high precision, considerably more so than the baseline method of a linear classifier. The recall rate of POEM is lower than that of the linear classifier, but that is expected as the linear classifier recovers all proximal genes with a distance of ≤ 500 bp. This means that the recall is high, but the number of false positives is also high, as indicated by the lower precision when compared to the CNN (Table 2, 69.84).

When recovering operons from metagenomes (Table 2), POEM’s results depend heavily upon the choice of gene-calling software and metagenome assembly. POEM outperforms the linear baseline method indicating that higher quality assemblies and longer reads will lead to a higher overall accuracy in POEM’s performance relative to the linear classifier. Furthermore, when recovering full operons, POEM’s CNN outperforms the linear classifier. The recovery overall is around 39% (1025 out of 2589), but it is considerably higher than that of the linear classifier (469/1,025).

the chief utility of POEM lies in identifying the functions carried out by the predicted operons in a metagenome. To that end, we introduced the core operon, identified by counting proximal predicted inter-operonic gene pairs in assembled contigs, and concatenating them using guilt-by-association. (Figure 5). The most frequent functions in the operons containing a large number of orthologous genes will be represented in the core operon. A high overlap in the count of functions (as COGs) between the core operons and the true operons indicates that while not all genes in an operon can be recovered in a metagenome, the basic functionality enabled by core operons can be recovered. The high precision and recall values shown in Table 4 indicate the the use of core operons can indeed inform us of those functions that are carried out by operons in a metagenome. In providing a characterization of core operons and their functions, POEM allows the annotation of a metagenome beyond the simple assignment of functions to genes, but to incorporate a level of annotation than includes an element of gene structure which is crucial in understanding bacterial function.

In sum, POEM is a novel and highly useful addition to the arsenal of tools helping us to better understand the functionality of metagenome, and is distinguished by offering a structural view of the metagenome, rather than a bag-of-genes-and-functions that most tools offer.

## Methods

An overview of the POEM pipeline is shown in Fig. 3. The heart of the pipeline lie the Operon identification and operon core structure that POEM performs. The other steps are performed with third-party tools, and are modular. Below we elaborate upon the various stages in the POEM pipeline.

**Figure 3.**
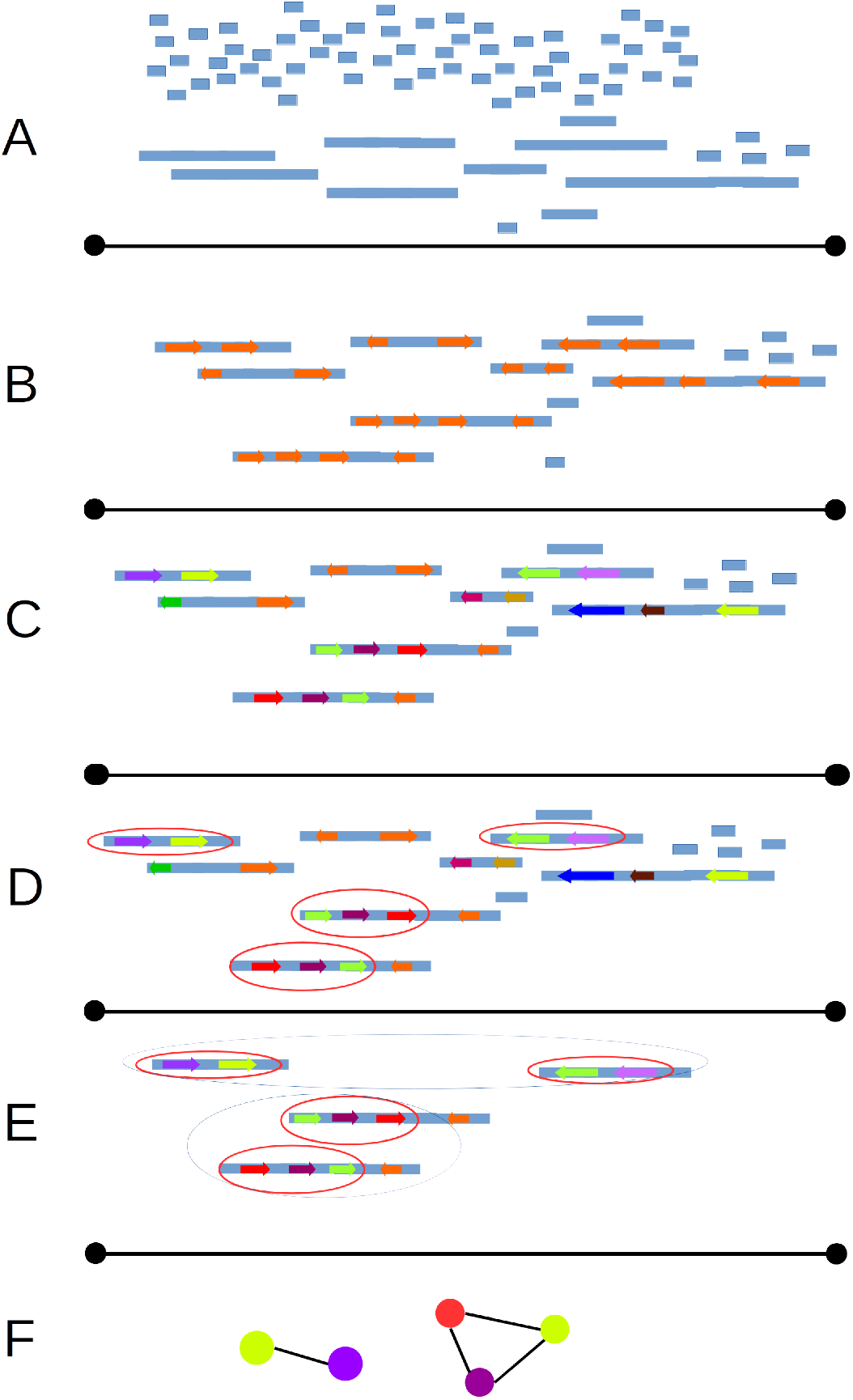
A flowcart of the POEM pipeline. A: assembly; B: Gene calling; C: similarity clustering; D: identify intra-operonic genes; E: identify core operons; F: graph-based visualization

### Metagenome Assembly

POEM uses, as default, the IDBA-UD *de-novo* assembler, but the user may supply an alternative assembler. Short read assemblers are usually based on De Bruijn graphs and are sensitive to the sequencing depth, repetitive regions, and sequencing errors [24]. For clonal bacteria, this assembly algorithm works relatively because it is easy to estimate the sequencing depth and the bacterial genomes are often compact and have few repetitive regions. However, in metagenomes it is hard to estimate the amount of sequence data that are needed for good functional coverage, and the genomes from closely related species may contain many highly conserved genes which may be interpreted as repetitive regions. Although *de novo* assemblers for metagenomes are still at an early stage [40], there are several tools developed for this task including MetaVelvet-SL, IDBA-UD,and Megahit [1, 22, 27, 30, 32]. In this study we also compare the effect these assemblers have on the accuracy of POEM.

### Gene Prediction

We chose to use an *ab-initio* method for gene calling, as opposed to calling by sequence similarity. First, because *ab-initio* gene calling is faster in bacterial and archaeal genomes, with little accuracy sacrificed: the predicted accuracy of some methods can reach 98% [16, 17, 43, 48]. Second, metagenomic data contain many genes with no similarity to known genes, so using a homology based method may result in a large number of open reading frames (ORFs) that are not predicted as such (false negatives). Several gene prediction tools have been developed or optimized for metagenomic data, including Glimmer-MG, Metagene, Metagenemark, Prokka, Prodigal, and Orphelia [14, 17, 20, 28, 36, 48]. POEM uses Metagenemark or Prokka to predict genes. As in the contig assembly stage, this part can be modified by the user.

### Removing ORF Redundancies

Once ORFs are identified, we remove redundant ORFs with an ID of >98% using CD-HIT [12, 23]. The assumption is that genes with a very high sequence ID were taken from the same species or highly similar strains and are therefore redundant information.

### Gene Function Annotation

While there are many ways to annotate gene function [34], a fast and acceptably accurate way to do so typically employs sequence similarity matching against a reliable functionally annotated sequence database. Here we used the COG database as a reference. POEM uses both BLAST and DIAMOND [4], which trades off speed and sensitivity. The functional assignment is done by choosing the top hit in COG above the e-value threshold (*Evalue* = 10^*−*3^).

### Operon Prediction

At the core of POEM lies a novel method we developed for predicting operons. POEM predicts if any given pair of adjacent genes are intra-operonic by classifying intergenic regions into intra- or extra-operonic. Thus, the operon prediction problem is cast as a binary classification problem.

POEM’s operon prediction method goes through the following steps. First, the intergenic DNA sequences of 4,425 operonic and 2,097 non-operonic adjacent genes were extracted from Operon DataBase v2 [29]. The intergenic regions are represented as a *k*-mer-position matrix (KPM, Figure 4). Two-thirds of the data were used for training a Convolutional Neural Network (CNN) based binary classification model and the remaining 1/3 of the data were used as the test set. We used a CNN model from the Keras package (v1.2.0) to train the classification model [7]. Since the CNN only accepts a fixed size matrix, we convert the KPM to a fixed size matrix by truncating the middle columns or adding all zero columns to the middle of the matrix. Trial-and-error has shown that *k* = 3 produced the best accuracy (Supplementary Figure S1).

**Figure 4.**
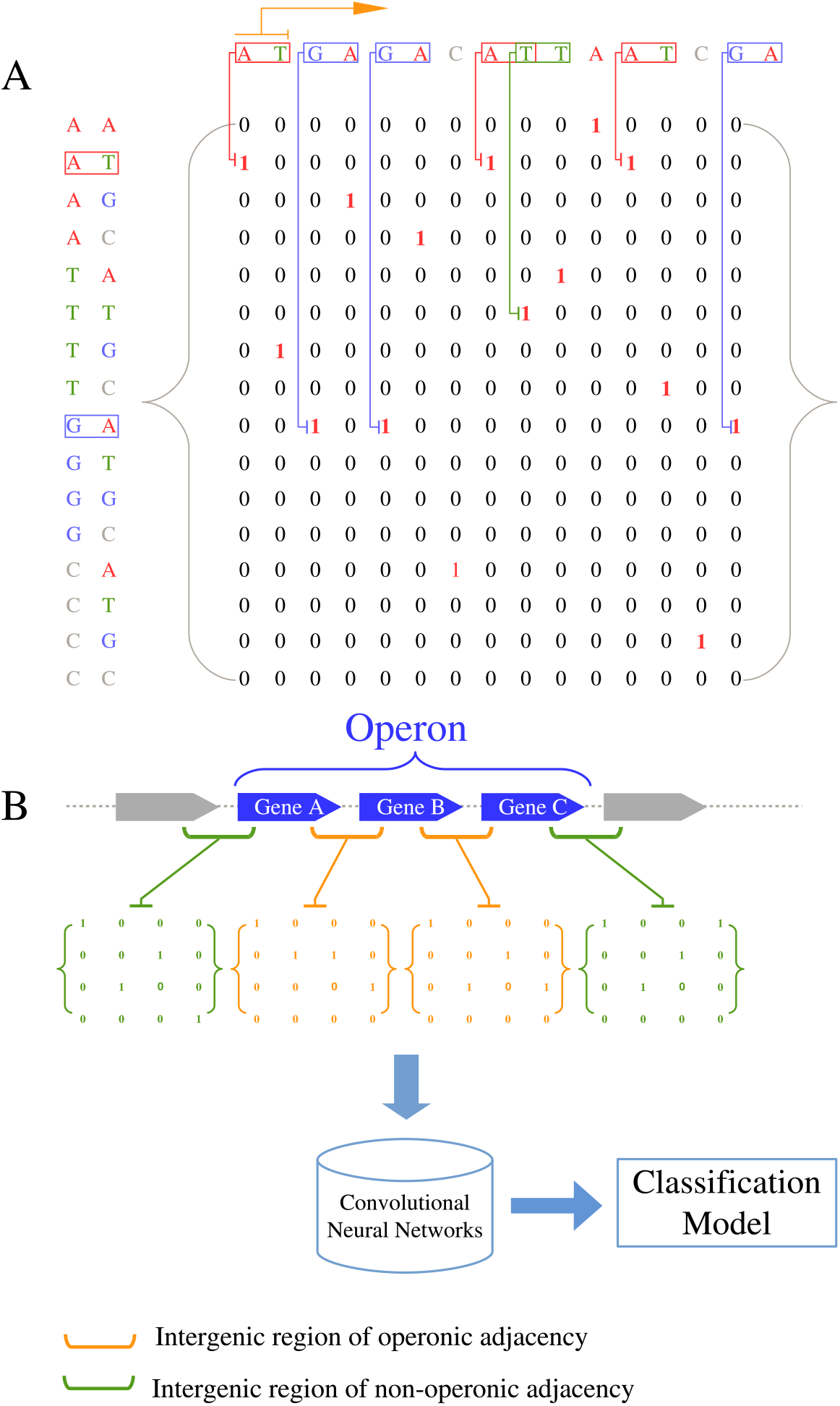
**A.** Construction of a *k*-mer-position matrix, shown with a 2-mer example (POEM uses 3-mer). Each row is a *k*-mer and the column number stands for a position in the sequence. If a specific *k*-mer appears in the sequence, the corresponding cell of the KPM is set to 1, otherwise, 0; **B.** training and building an CNN based classification model from intergenic of operonic and non-operonic adjacency.

**Figure 5.**
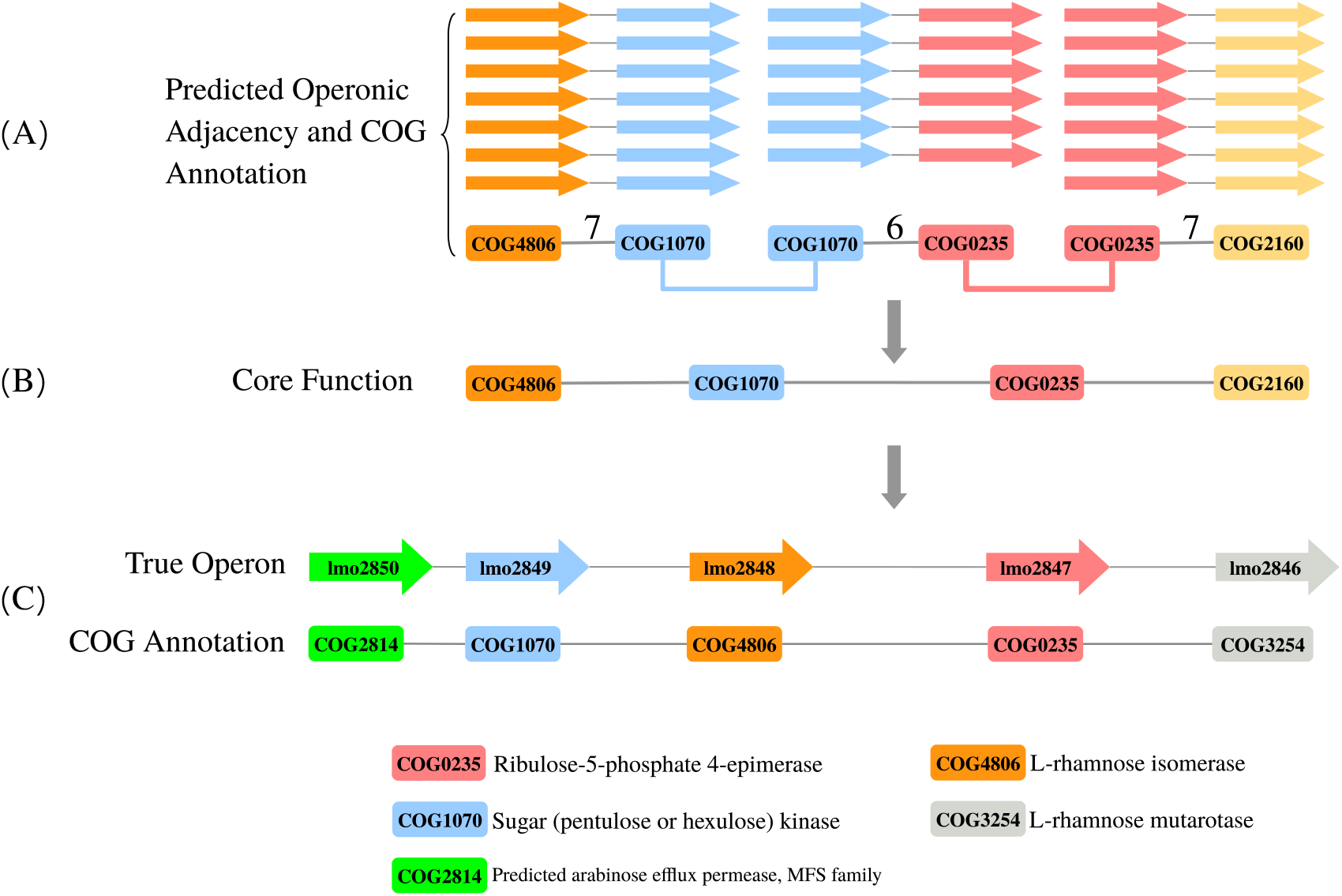
Identifying Core Operons. **A:** find orthologous COG-annotated proximal gene pairs and concatenate them using guilt-by-association. **B:** The resulting graph shows the *core function* (four different COG IDs) **C:** Find the most similar operon in the dataset of gold standards and its corresponding GO annotations. In this example, there are 3 true positives (COG4806, COG1070, and COG0235), 1 false positive (COG2160), and 2 false negatives (COG2814 & COG3254). Precision is therefore 0.75 and recall is 0.6

To show the CNN’s utility, we compared its performance to a simple baseline predictor. The baseline linear classifier works as follows: if two genes on the same strand have an intergenic distance < 500 nt, then their adjacency is classified as within the same operon (operonic). A larger distance would classify them as non-operonic. The predicted operonic adjacent genes were then connected to form a full operon prediction.

### Identifying Core Operons

To characterize operons in metagenomes, we introduce the concept of *core operons*. Core operons are weighted-edge undirected graphs that capture information about predicted orthologous operons or fractions of operons in the metagenome. Each node is a set of orthologous genes that are all annotated by at least one common COG term. An edge is drawn between two nodes if they are determined to be an intra-operonic pair. The weight of the edge is determined by the frequency of the adjacency of the intra-operon adjacent genes. To determine how well a core operon captures the real operons in a metagenome, we ran a precision-recall analysis using the operons in the simulated database as our standard-of-truth, see Figure 5. Here, precision is the number of correctly predicted intra-operonic genes (true positives) divided by the number of all predictions (true positive and false positive predictions). Recall is the number of correctly predicted intra-operonic genes divided by the all real intra-operonic genes. Finally, POEM produces a file that can then be used by Cytoscape [37] to visualize the core operons.

## Availability of source code and requirements

The software and related information are listed below:

**Project Name:** POEM

**Project Home Page:** https://github.com/Rinoahu/POEM_py3k

**Operating System(s):** POEM was tested on GNU/Linux distribution Ubuntu 16.04 64-bit, but we expect POEM to work on most Unix-like systems.

**Programming Language:** Python

**Other Requirements:** Python 3.7 and Conda

**License:** GPLv3

## Acknowledgments

Will be provided upon acceptance

## References

1. Afiahayati, Kengo Sato, and Yasubumi Sakakibara. MetaVelvet-SL: An extension of the Velvet assembler to a de novo metagenomic assembler utilizing supervised learning. DNA Res., 22(1):69–77, 2015.

2. Dennis A. Benson, Karen Clark, Ilene Karsch-Mizrachi, David J. Lipman, James Ostell, and Eric W. Sayers. GenBank. Nucleic Acids Res., 2015.

3. J. Bockhorst, M. Craven, D. Page, J. Shavlik, and J. Glasner. A Bayesian network approach to operon prediction. Bioinformatics, 19(10):1227–1235, jul 2003.

4. Benjamin Buchfink, Chao Xie, and Daniel H Huson. Fast and sensitive protein alignment using DIAMOND. Nat. Methods, 12(1):59–60, nov 2014.

5. X Chen, Z Su, P Dam, B Palenik, Y Xu, and T Jiang. Operon prediction by comparative genomics: an application to the Synechococcus sp. WH8102 genome. Nucleic Acids Res., 32(7):2147–57, 2004.

6. Joshua L Cherry. Genome size and operon content. Journal of theoretical biology, 221(3):401–410, 2003.

7. François Chollet. Keras: Deep Learning library for Theano and TensorFlow, 2015.

8. Li-Yeh Chuang, Jui-Hung Tsai, and Cheng-Hong Yang. Binary particle swarm optimization for operon prediction. Nucleic Acids Res., 38(12):e128, jul 2010.

9. Li-Yeh Chuang, Cheng-Huei Yang, Jui-Hung Tsai, and Cheng-Hong Yang. Operon Prediction Using Chaos Embedded Particle Swarm Optimization. IEEE/ACM Trans. Comput. Biol. Bioinforma., 10(5):1299–1309, sep 2013.

10. Mark Craven, David Page, Jude Shavlik, Joseph Bockhorst, and Jeremy Glasner. A Probabilistic Learning Approach to Whole-Genome Operon Prediction. Proc Int Conf Intell Syst Mol Biol, 8:116–27, 2000.

11. Phuongan Dam, Victor Olman, Kyle Harris, Zhengchang Su, and Ying Xu. Operon prediction using both genome-specific and general genomic information. Nucleic Acids Res., 35(1):288–298, 2007.

12. Limin Fu, Beifang Niu, Zhengwei Zhu, Sitao Wu, and Weizhong Li. CD-HIT: Accelerated for clustering the next-generation sequencing data. Bioinformatics, 28(23):3150–3152, 2012.

13. Socorro Gama-Castro, Heladia Salgado, Alberto Santos-Zavaleta, Daniela Ledezma-Tejeida, Luis Muñiz-Rascado, Jair Santiago García-Sotelo, Kevin Alquicira-Hernández, Irma Martínez-Flores, Lucia Pannier, Jaime Abraham Castro-Mondragón, et al. Regulondb version 9.0: high-level integration of gene regulation, coexpression, motif clustering and beyond. Nucleic acids research, 44(D1):D133–D143, 2016.

14. Katharina J. Hoff, Thomas Lingner, Peter Meinicke, and Maike Tech. Orphelia: Predicting genes in metagenomic sequencing reads. Nucleic Acids Res., 37(SUPPL. 2), 2009.

15. Weichun Huang, Leping Li, Jason R. Myers, and Gabor T. Marth. ART: A next-generation sequencing read simulator. Bioinformatics, 28(4):593–594, 2012.

16. Doug Hyatt, Gwo-Liang Chen, Philip F Locascio, Miriam L Land, Frank W Larimer, and Loren J Hauser. Prodigal: prokaryotic gene recognition and translation initiation site identification. BMC Bioinformatics, 11:119, 2010.

17. Doug Hyatt, Philip F. Locascio, Loren J. Hauser, and Edward C. Uberbacher. Gene and translation initiation site prediction in metagenomic sequences. Bioinformatics, 28(17):2223–2230, 2012.

18. E. Jacob, R. Sasikumar, and K. N. R. Nair. A fuzzy guided genetic algorithm for operon prediction. Bioinformatics, 21(8):1403–1407, apr 2005.

19. Minoru Kanehisa and Susumu Goto. KEGG: Kyoto Encyclopedia of Genes and Genomes. Nucleic Acids Research, 28(1):27–30, January 2000.

20. David R. Kelley, Bo Liu, Arthur L. Delcher, Mihai Pop, and Steven L. Salzberg. Gene prediction with Glimmer for metagenomic sequences augmented by classification and clustering. Nucleic Acids Res., 40(1), 2012.

21. Rasko Leinonen, Ruth Akhtar, Ewan Birney, Lawrence Bower, Ana Cerdeno-Tárraga, Ying Cheng, Iain Cleland, Nadeem Faruque, Neil Goodgame, Richard Gibson, Gemma Hoad, Mikyung Jang, Nima Pakseresht, Sheila Plaister, Rajesh Radhakrishnan, Kethi Reddy, Siamak Sobhany, Petra Ten Hoopen, Robert Vaughan, Vadim Zalunin, and Guy Cochrane. The European nucleotide archive. Nucleic Acids Res., 39(SUPPL. 1), 2011.

22. Dinghua Li, Chi Man Liu, Ruibang Luo, Kunihiko Sadakane, and Tak Wah Lam. MEGAHIT: An ultra-fast single-node solution for large and complex metagenomics assembly via succinct de Bruijn graph. Bioinformatics, 31(10):1674–1676, 2014.

23. Weizhong Li and Adam Godzik. Cd-hit: A fast program for clustering and comparing large sets of protein or nucleotide sequences. Bioinformatics, 22(13):1658–1659, 2006.

24. Zhenyu Li, Yanxiang Chen, Desheng Mu, Jianying Yuan, Yujian Shi, Hao Zhang, Jun Gan, Nan Li, Xuesong Hu, Binghang Liu, Bicheng Yang, and Wei Fan. Comparison of the two major classes of assembly algorithms: overlap-layout-consensus and de-bruijn-graph. Brief. Funct. Genomics, 11(1):25–37, jan 2012.

25. Fenglou Mao, Phuongan Dam, Jacky Chou, Victor Olman, and Ying Xu. Door: a database for prokaryotic operons. Nucleic acids research, 37(suppl 1):D459–D463, 2009.

26. Gabriel Moreno-Hagelsieb and Julio Collado-Vides. A powerful non-homology method for the prediction of operons in prokaryotes. Bioinformatics, 18 Suppl 1:S329–36, 2002.

27. Toshiaki Namiki, Tsuyoshi Hachiya, Hideaki Tanaka, and Yasubumi Sakakibara. MetaVelvet: An extension of Velvet assembler to de novo metagenome assembly from short sequence reads. Nucleic Acids Res., 40(20), 2012.

28. Hideki Noguchi, Jungho Park, and Toshihisa Takagi. MetaGene: Prokaryotic gene finding from environmental genome shotgun sequences. Nucleic Acids Res., 34(19):5623–5630, 2006.

29. Shujiro Okuda and Akiyasu C Yoshizawa. Odb: a database for operon organizations, 2011 update. Nucleic acids research, 39(suppl 1):D552–D555, 2011.

30. Anastasis Oulas, Christina Pavloudi, Paraskevi Polymenakou, Georgios A Pavlopoulos, Nikolas Papanikolaou, Georgios Kotoulas, Christos Arvanitidis, and Ioannis Iliopoulos. Metagenomics: tools and insights for analyzing next-generation sequencing data derived from biodiversity studies. Bioinform. Biol. Insights, 9:75–88, 2015.

31. R. Overbeek, M. Fonstein, M. D’Souza, G. D. Pusch, and N. Maltsev. The use of gene clusters to infer functional coupling. Proceedings of the National Academy of Sciences of the United States of America, 96(6):2896–2901, March 1999.

32. Yu Peng, Henry C M Leung, S. M. Yiu, and Francis Y L Chin. IDBA-UD: A de novo assembler for single-cell and metagenomic sequencing data with highly uneven depth. Bioinformatics, 28(11):1420–1428, 2012.

33. Mihaela Pertea, Kunmi Ayanbule, Megan Smedinghoff, and Steven L Salzberg. Operondb: a comprehensive database of predicted operons in microbial genomes. Nucleic acids research, 37(suppl 1):D479–D482, 2009.

34. Predrag Radivojac, Wyatt T Clark, Tal Ronnen Oron, Alexandra M Schnoes, Tobias Wittkop, Artem Sokolov, Kiley Graim, Christopher Funk, Karin Verspoor, Asa Ben-Hur, et al. A large-scale evaluation of computational protein function prediction. Nature methods, 10(3):221, 2013.

35. C. Sabatti, Lars Rohlin, Min-Kyu Oh, and James C. Liao. Co-expression pattern from DNA microarray experiments as a tool for operon prediction. Nucleic Acids Res., 30(13):2886–2893, jul 2002.

36. Torsten Seemann. Prokka: rapid prokaryotic genome annotation. Bioinformatics, 30(14):2068–2069, 03 2014.

37. Paul Shannon, Andrew Markiel, Owen Ozier, Nitin S. Baliga, Jonathan T. Wang, Daniel Ramage, Nada Amin, Beno Schwikowski, and Trey Ideker. Cytoscape: A software Environment for integrated models of biomolecular interaction networks. Genome Res., 13(11):2498–2504, 2003.

38. Blanca Taboada, Ricardo Ciria, Cristian E Martinez-Guerrero, and Enrique Merino. Proopdb: prokaryotic operon database. Nucleic acids research, 40(D1):D627–D631, 2012.

39. Blanca Taboada, Cristina Verde, and Enrique Merino. High accuracy operon prediction method based on STRING database scores. Nucleic Acids Res., 38(12):e130, jul 2010.

40. Torsten Thomas, Jack Gilbert, and Folker Meyer. Metagenomics - a guide from sampling to data analysis. Microbial Informatics and Experimentation, 2(1):3, February 2012.

41. Gregory Vey and Trevor C. Charles. Metaprox: the database of metagenomic proximons. Database: the journal of biological databases and curation, 2014, 2014.

42. Gregory Vey and Gabriel Moreno-Hagelsieb. Metagenomic annotation networks: Construction and applications. PLoS One, 7(8), 2012.

43. Zhuo Wang, Yazhu Chen, and Yixue Li. A brief review of computational gene prediction methods. Genomics, proteomics Bioinforma. / Beijing Genomics Inst., 2(4):216–21, 2004.

44. Yuri I Wolf, Igor B Rogozin, Alexey S Kondrashov, and Eugene V Koonin. Genome alignment, evolution of prokaryotic genome organization, and prediction of gene function using genomic context. Genome research, 11(3):356–372, 2001.

45. John C. Wooley, Adam Godzik, and Iddo Friedberg. A Primer on Metagenomics. PLoS Comput. Biol., 6(2):e1000667, feb 2010.

46. Syed Shujaat Ali S. Zaidi and Xuegong Zhang. Computational operon prediction in whole-genomes and metagenomes. Briefings in functional genomics, September 2016.

47. Daniel R. Zerbino and Ewan Birney. Velvet: Algorithms for de novo short read assembly using de Bruijn graphs. Genome Res., 18(5):821–829, 2008.

48. Wenhan Zhu, Alexandre Lomsadze, and Mark Borodovsky. Ab initio gene identification in metagenomic sequences. Nucleic Acids Res., 38(12), 2010.

