## Supplementary file for "Identifying Core Operons in Metagenomic Data"

### Supplementary materials for: "Identifying Core Operons in Metagenomic Data"

Xiao Hu and Iddo Friedberg

January 28, 2020

#### Third party software used

**Simulated metagenome** We use ART(version "GSM.04.18.2016") [1] to generate simulated Illumina paired-reads for each genome. The command line is:

```
$art_illumina -ss HS25 -sam -i input -p -l 150 -f X -m 200 -s 30
-o output.
```

Read length was 150 nt. The mean and standard deviation of fragment length is 200 nt and 30, respectively. Parameter -f X is the fold coverage of the reads for a specific genome. The name, gi number and fold coverage for each specific genome are list in table S1. We then put all the reads in a single file.

**Assembly with MegaHIT:**

```
$megahit --presets meta-sensitive -1 left.fq -2 right.fq -o megahit_asm
```

**Assembly with IDBA:**

```
$idba_ud -r reads.fa -o idba_output --num_threads 16
-o output
```

**Assembly with Velvet:**

```
$velveth asm_51 51 -fastq -shortPaired -separate left.fq right.fq
$velvetg asm_51 -exp_cov auto -ins_length 200 -scaffolding no -read_trkg yes
```

**Gene clustering:** CD-HIT was used as follows:

```
$cd-hit -c .70 -aL .85 -M 12000 -T 0 -d 268435456 -i input -o output
```

**Gene prediction:**

```
$gmhmmp -A output -p 0 -f G -m model input
```

The HMM model file with gene finding parameters used (-m) from QUAST v2.3 [2].

**Precision, recall and F1 score of operon prediction under different *k*-mer lengths.**

<table>
<tr><th>Species</th><th>GI</th><th>Abbre</th><th>-f X</th></tr>
<tr><td>Caenorhabditis elegans mitochondrion, complete genome</td><td>gi|5834884|ref|NC_001328.1|</td><td>cel</td><td>5X</td></tr>
<tr><td>Prochlorococcus marinus subsp. marinus str. CCMP1375 complete genome</td><td>gi|33239452|ref|NC_005042.1|</td><td>pma</td><td>5X</td></tr>
<tr><td>Porphyromonas gingivalis W83, complete genome</td><td>gi|34539880|ref|NC_002950.2|</td><td>pgi</td><td>5X</td></tr>
<tr><td>Haemophilus influenzae Rd KW20 chromosome, complete genome</td><td>gi|16271976|ref|NC_000907.1|</td><td>hin</td><td>8X</td></tr>
<tr><td>Nitrosomonas europaea ATCC 19718, complete genome</td><td>gi|30248031|ref|NC_004757.1|</td><td>neu</td><td>5X</td></tr>
<tr><td>Clostridium acetobutylicum ATCC 824 chromosome, complete genome</td><td>gi|15893298|ref|NC_003030.1|</td><td>cac</td><td>5X</td></tr>
<tr><td>Mycobacterium bovis subsp. bovis AF2122/97 complete genome</td><td>gi|31742509|emb|BX248333.1|</td><td>mbo</td><td>5X</td></tr>
<tr><td>Haemophilus ducreyi strain 35000HP complete genome</td><td>gi|33151282|ref|NC_002940.2|</td><td>hdu</td><td>5X</td></tr>
<tr><td>Yersinia pestis CO92, complete genome</td><td>gi|755429805|gb|CP009973.1|</td><td>ype</td><td>5X</td></tr>
<tr><td>Neisseria meningitidis MC58 chromosome, complete genome</td><td>gi|77358697|ref|NC_003112.2|</td><td>nme</td><td>7X</td></tr>
<tr><td>Corynebacterium glutamicum ATCC 13032, IS fingerprint type 4-5, complete genome</td><td>gi|62388892|ref|NC_006958.1|</td><td>cgl</td><td>9X</td></tr>
<tr><td>Yersinia pseudotuberculosis IP 32953, complete genome</td><td>gi|756143888|ref|NZ_CP009712.1|</td><td>yps</td><td>7X</td></tr>
<tr><td>Pyrococcus furiosus DSM 3638, complete genome</td><td>gi|18976372|ref|NC_003413.1|</td><td>pfu</td><td>5X</td></tr>
<tr><td>Synechocystis sp. PCC 6803 substrain GT-G, complete genome</td><td>gi|939195038|gb|CP012832.1|</td><td>syn</td><td>13X</td></tr>
<tr><td>Ciona intestinalis B CG-2006 mitochondrion, complete genome</td><td>gi|387935481|ref|NC_017929.1|</td><td>cin</td><td>5X</td></tr>
<tr><td>Drosophila melanogaster isolate ZIM mitochondrion, complete genome</td><td>gi|848113863|gb|KP843854.1|</td><td>dme</td><td>69X</td></tr>
<tr><td>Bacillus subtilis subsp. subtilis str. 168, complete genome</td><td>gi|749168884|ref|NZ_CP010052.1|</td><td>bsu</td><td>5X</td></tr>
<tr><td>Salmonella enterica subsp. enterica serovar Typhi Ty2, complete genome</td><td>gi|29140543|ref|NC_004631.1|</td><td>stt</td><td>5X</td></tr>
<tr><td>Rhodopseudomonas palustris CGA009 complete genome</td><td>gi|39933080|ref|NC_005296.1|</td><td>rpa</td><td>5X</td></tr>
<tr><td>Campylobacter jejuni subsp. jejuni NCTC 11168-K12E5, complete genome</td><td>gi|752706567|ref|NZ_CP006685.1|</td><td>cje</td><td>6X</td></tr>
<tr><td>Nostoc sp. PCC 7120 DNA, complete genome</td><td>gi|17227497|ref|NC_003272.1|</td><td>ana</td><td>5X</td></tr>
<tr><td>Staphylococcus aureus subsp. aureus N315 DNA, complete genome</td><td>gi|29165615|ref|NC_002745.2|</td><td>sau</td><td>18X</td></tr>
<tr><td>Sulfolobus solfataricus P2, complete genome</td><td>gi|15896971|ref|NC_002754.1|</td><td>sso</td><td>8X</td></tr>
<tr><td>Bradyrhizobium japonicum USDA 110 chromosome, complete genome</td><td>gi|27375111|ref|NC_004463.1|</td><td>bja</td><td>9X</td></tr>
<tr><td>Clostridium perfringens str. 13 DNA, complete genome</td><td>gi|18308982|ref|NC_003366.1|</td><td>cpe</td><td>25X</td></tr>
<tr><td>Lactobacillus plantarum WCFS1, complete genome</td><td>gi|380031102|ref|NC_004567.2|</td><td>lpl</td><td>20X</td></tr>
<tr><td>Helicobacter pylori 26695, complete genome</td><td>gi|410024832|ref|NC_018939.1|</td><td>hpy</td><td>11X</td></tr>
<tr><td>Salmonella enterica subsp. enterica serovar Typhimurium str. 14028S, complete genome</td><td>gi|267991652|gb|CP001363.1|</td><td>stm</td><td>15X</td></tr>
<tr><td>Acinetobacter sp. ADP1 complete genome</td><td>gi|50083297|ref|NC_005966.1|</td><td>aci</td><td>31X</td></tr>
<tr><td>Mycoplasma pneumoniae M129 chromosome, complete genome</td><td>gi|13507739|ref|NC_000912.1|</td><td>mpn</td><td>69X</td></tr>
<tr><td>Photorhabdus luminescens subsp. laumondii TTO1 complete genome</td><td>gi|37524032|ref|NC_005126.1|</td><td>plu</td><td>5X</td></tr>
<tr><td>Borrelia burgdorferi B31 chromosome, complete genome</td><td>gi|15594346|ref|NC_001318.1|</td><td>bbu</td><td>6X</td></tr>
<tr><td>Treponema denticola ATCC 35405 chromosome, complete genome</td><td>gi|42516522|ref|NC_002967.9|</td><td>tde</td><td>14X</td></tr>
<tr><td>Desulfovibrio vulgaris str. Hildenborough chromosome, complete genome</td><td>gi|46562128|ref|NC_002937.3|</td><td>dvu</td><td>5X</td></tr>
<tr><td>Escherichia coli str. K-12 substr. MG1655, complete genome</td><td>gi|985000614|gb|CP014225.1|</td><td>eco</td><td>5X</td></tr>
<tr><td>Lactococcus lactis subsp. lactis Il1403 chromosome, complete genome</td><td>gi|15671982|ref|NC_002662.1|</td><td>lla</td><td>12X</td></tr>
<tr><td>Pseudomonas aeruginosa PAO1 chromosome, complete genome</td><td>gi|110645304|ref|NC_002516.2|</td><td>pae</td><td>21X</td></tr>
<tr><td>Lactobacillus johnsonii NCC 533, complete genome</td><td>gi|42518084|ref|NC_005362.1|</td><td>ljo</td><td>5X</td></tr>
<tr><td>Mycobacterium tuberculosis H37Rv, complete genome</td><td>gi|752689705|ref|NZ_CP009480.1|</td><td>mtu</td><td>22X</td></tr>
<tr><td>Methanothermobacter thermautotrophicus str. Delta H, complete genome</td><td>gi|15678031|ref|NC_000916.1|</td><td>mth</td><td>66X</td></tr>
<tr><td>Aquifex aeolicus VF5 chromosome, complete genome</td><td>gi|15282445|ref|NC_000918.1|</td><td>aae</td><td>80X</td></tr>
<tr><td>Bordetella pertussis Tohama I chromosome, complete genome</td><td>gi|33591275|ref|NC_002929.2|</td><td>bpe</td><td>16X</td></tr>
<tr><td>Streptomyces coelicolor A3(2) complete genome</td><td>gi|30407153|emb|AL645882.2|</td><td>sco</td><td>16X</td></tr>
<tr><td>Staphylococcus aureus subsp. aureus strain MRSA252, complete genome</td><td>gi|49482253|ref|NC_002952.2|</td><td>sar</td><td>100X</td></tr>
<tr><td>Listeria monocytogenes EGD-e, complete genome</td><td>gi|30407125|emb|AL591824.1|</td><td>lmo</td><td>104X</td></tr>
<tr><td>Yersinia pestis KIM10+, complete genome</td><td>gi|22123922|ref|NC_004088.1|</td><td>ypk</td><td>91X</td></tr>
<tr><td>Bacillus cereus ATCC 14579 chromosome, complete genome</td><td>gi|30018278|ref|NC_004722.1|</td><td>bce</td><td>72X</td></tr>
<tr><td>Bordetella bronchiseptica strain RB50, complete genome</td><td>gi|33598993|ref|NC_002927.3|</td><td>bbr</td><td>82X</td></tr>
</table>

Table S1: Genomes used to generate simulated illumina reads. **Abbre:** Abbreviation used by Operon Database [3]; **-f x:** Fold coverage of simulated reads for corresponding genome.

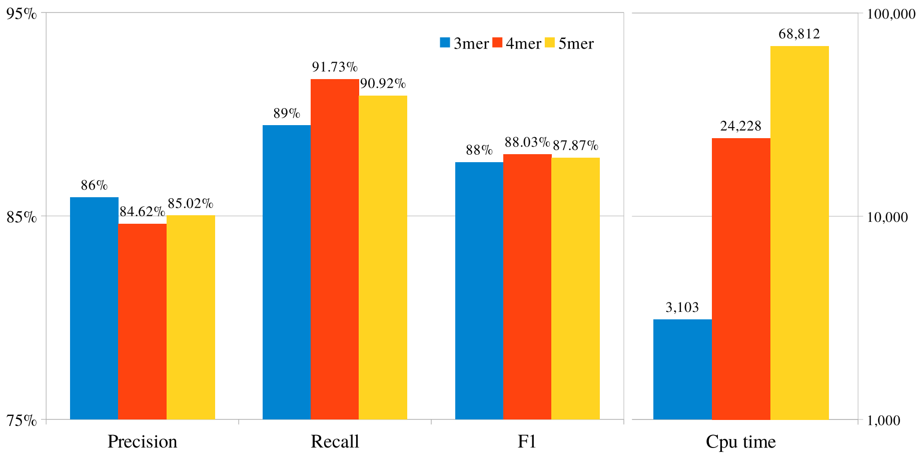


Figure S1: Precision, recall and F1 score of operon prediction under different $k$-mer lengths. The performance of different kmers are similar. Using 3-mer is considerably faster than 4-mer or 5- mer.

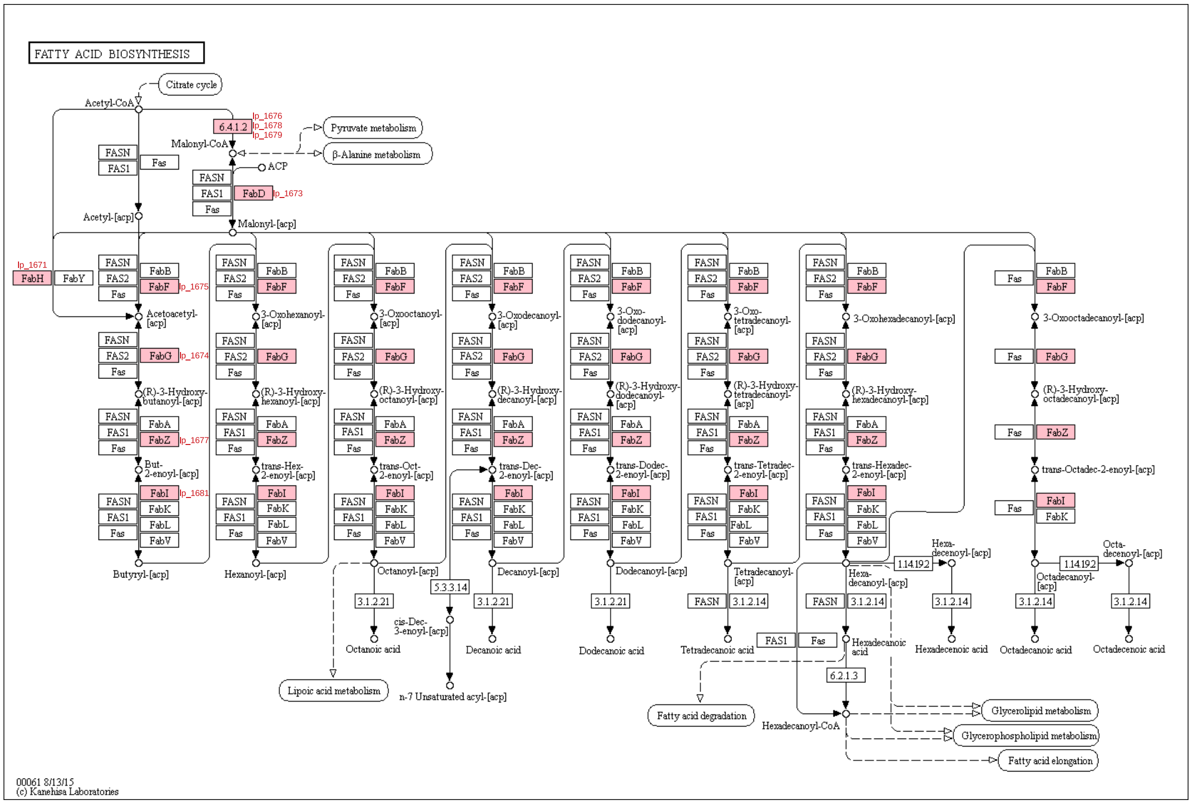


Figure S2: Mapping predicted operonic genes in *Lactobacillus plantarum* WCFS1 to KEGG pathway. The gene in pink box is the predicted operonic gene.

| **Locus** | **start** | **end** | **strand** | **Gene** | **COG** | **KEGG** | **Function** |
|---|---|---|---|---|---|---|---|
| | | | | ***Bacillus cereus ATCC 10987*** | | | |
| BCE_3896 | 3644360 | 3644953 | - | fapR | COG2050 | - | Transcription factor FapR |
| **BCE_3895** | **3643371** | **3644363** | **-** | **plsX** | **COG0416** | **K03621** | **fatty acid/phospholipid synthesis protein PlsX** |
| **BCE_3894** | **3642412** | **3643356** | **-** | **fabD** | **COG0331** | **K00645** | **malonyl CoA-acyl carrier protein transacylase** |
| **BCE_3893** | **3641672** | **3642412** | **-** | **fabG** | **COG1028** | **K00059** | **3-oxoacyl-(acyl-carrier-protein) reductase** |
| **BCE_3892** | **3641369** | **3641602** | **-** | **acpA** | **COG0236** | **K02078** | **acyl carrier protein** |
| BCE_3891 | 3640573 | 3641310 | - | rncS | COG0571 | K03685 | ribonuclease III |
| | | | | ***Megamonas funiformis YIT 11815*** | | | |
| EHR36953 | 11106 | 12110 | - | | unknown | K00648 | 3-oxoacyl-[acyl-carrier-protein] synthase 3 |
| EHR36952 | 10155 | 11102 | - | | COG2070 | K02371 | putative enoyl-(acyl-carrier-protein) reductase |
| **EHR36951** | **9168** | **10112** | **-** | | **COG0331** | **K00645** | **malonyl CoA-acyl carrier protein transacylase** |
| **EHR36950** | **8423** | **9166** | **-** | | **COG1028** | **K00059** | **3-oxoacyl-[acyl-carrier-protein] reductase** |
| **EHR36949** | **8148** | **8378** | **-** | | **COG0236** | **K02078** | **acyl carrier protein** |
| EHR36948 | 7116 | 8060 | - | | COG2070 | K02371 | hypothetical protein |
| **EHR36947** | **5858** | **7099** | **-** | | **COG0304** | **-** | **beta-ketoacyl-acyl-carrier-protein synthase II** |
| EHR36946 | 5133 | 5843 | - | | COG0571 | K03685 | ribonuclease III |
| | | | | ***Ruminococcus sp. AM09-18-1*** | | | |
| RGF28942 | 23266 | 24213 | + | | unkown | K00648 | ketoacyl-ACP synthase III |
| **RGF29171** | **24237** | **24461** | **+** | | **COG0236** | **K02078** | **acyl carrier protein** |
| RGF28943 | 24559 | 25497 | + | fabK | COG2070 | K02371 | enoyl-[acyl-carrier-protein] reductase FabK |
| **RGF28944** | **25490** | **26410** | **+** | **fabD** | **COG0331** | **K00645** | **[acyl-carrier-protein] S-malonyltransferase** |
| **RGF28945** | **26404** | **27153** | **+** | **fabG** | **COG1028** | **K00059** | **3-oxoacyl-[acyl-carrier-protein] reductase** |
| **RGF28946** | **27173** | **28411** | **+** | **fabF** | **COG0304** | **K09458** | **beta-ketoacyl-[acyl-carrier-protein] synthase II** |
| RGF28947 | 28424 | 28918 | + | accB | unkown | K02160 | acetyl-CoA carboxylase biotin carboxyl carrier |
| RGF28948 | 28934 | 29365 | + | fabZ | COG0764 | K02372 | 3-hydroxyacyl-[acyl-carrier-protein] dehydratase FabZ |
| RGF28949 | 29386 | 30735 | + | | COG0439 | K01961 | acetyl-CoA carboxylase biotin carboxylase subunit |
| RGF28950 | 30722 | 32440 | + | | COG0825 | K01962 | acetyl-CoA carboxylase carboxyltransferase subunit beta |
| | | | | ***Lactobacillus plantarum WCFS1*** | | | |
| lp_1670 | 1522732 | 1523175 | + | fabZ | COG0764 | - | 3-hydroxyacyl-[acyl-carrier-protein] dehydratase FabZ |
| lp_1671 | 1523250 | 1524236 | + | fabH2 | COG0332 | K00648 | 3-oxoacyl-[acyl-carrier-protein] synthase 3 protein 2 |
| **lp_1672** | **1524293** | **1524541** | **+** | **acpA2** | **COG0236** | **K02078** | **acyl carrier protein** |
| **lp_1673** | **1524545** | **1525477** | **+** | **fabD** | **COG0331** | **K00645** | **[acyl-carrier protein] S-malonyltransferase** |
| **lp_1674** | **1525464** | **1526192** | **+** | **fabG1** | **COG1028** | **K00059** | **3-oxoacyl-ACP reductase** |
| **lp_1675** | **1526211** | **1527443** | **+** | **fabF** | **COG0304** | **K09458** | **3-oxoacyl-[acyl-carrier protein] synthase II** |
| lp_1676 | 1527440 | 1527901 | + | accB2 | COG0511 | K02160 | acetyl-CoA carboxylase, biotin carboxyl carrier protein |
| lp_1677 | 1527901 | 1528314 | + | fabZ2 | COG0764 | K02372 | (3R)-hydroxymyristoyl-[acyl carrier protein] dehydratase |
| lp_1678 | 1528329 | 1529717 | + | accC2 | COG0439 | K01961 | acetyl-CoA carboxylase, biotin carboxylase subunit |
| lp_1679 | 1529683 | 1530528 | + | accD2 | COG0777 | K01963 | acetyl-CoA carboxylase, carboxyl transferase subunit beta |
| lp_1680 | 1530521 | 1531291 | + | accA2 | COG0825 | K01962 | acetyl-CoA carboxylase, carboxyl transferase subunit alpha |
| lp_1681 | 1531312 | 1532070 | + | fabI | COG0623 | K00208 | enoyl-[acyl-carrier protein] reductase (NADH) |
| lp_1682 | 1532070 | 1532621 | + | | COG2091 | K06133 | phosphopantetheinyltransferase |

Table S2: Functional annotations for the predicted operonic genes from SRR2155174 data set. **COG**: Clusters of Orthologous Group identifier; **KEGG**: KEGG Orthology database; Bold locus names denote the genes found in the core function.

### References

[1] Weichun Huang, Leping Li, Jason R. Myers, and Gabor T. Marth. ART: A next-generation sequencing read simulator. *Bioinformatics*, 28(4):593–594, 2012.

[2] Alexey Gurevich, Vladislav Saveliev, Nikolay Vyahhi, and Glenn Tesler. QUAST: Quality assessment tool for genome assemblies. *Bioinformatics*, 29(8):1072–1075, 2013.

[3] Shujiro Okuda and Akiyasu C Yoshizawa. Odb: a database for operon organizations, 2011 update. *Nucleic acids research*, 39(suppl 1):D552–D555, 2011.
